# Alteration of epigenetic landscape by lamin A mutations: Hallmark of Dilated Cardiomyopathy

**DOI:** 10.1101/2020.05.01.071803

**Authors:** Avinanda Banerjee, Kaushik Sengupta

## Abstract

Mutations in lamin A have been reported to be associated with over 16 human diseases including dilated cardiomyopathy (DCM). We have focused on three such DCM causing mutants of lamin A which have to address the contribution of lamins in the pathogenesis of DCM at molecular level. We have elucidated the effect of these mutants for the first time on the epigenetic landscape of a myogenic fibroblast cell line C2C12. C2C12 cells expressing these mutant proteins exhibited alterations in some histone modification marks like H3K4me3, H3K9me3, H3K27me3, H3K36me3 and RNA Polymerase II activity compared to its wild type variants. This report paves the way for further studies involving epigenetic regulation in laminopathies which would be an important step in explaining the molecular mechanism and pathophysiology of the diseases like dilated cardiomyopathy.

**High Lights:**

1. lamin A K97E mutation predominantly alters H3K9me3 histone modifications landscape
2. lamin A K97E aggregates within nucleus also sequester the HP1γ
3. lamin A K97E mutation affects RNA polymerase II distribution pattern

## Introduction

Lamin proteins play an active role in nuclear homeostasis in most mammalian somatic cells by tethering and organizing chromatin as well as modulating epigenetic landscape [1,2]. Both A- and B-type lamins constituting the nuclear lamina dictates peripheral heterochromatin distribution as well as maintains nuclear integrity and rigidity [3]. In most mammalian cells A-type lamin manifests itself as a soluble nucleoplasmic form in addition to the lamina associated insoluble form whereas B-type lamins are largely insoluble and almost exclusively confined to the peripheral lamina [4]. Almost 500 mutations associated with lamin A have been shown to cause 16 different human disorders like Dilated Cardiomyopathy (DCM),[5]Emery-Dreifuss muscular dystrophy (AD-EDMD),[6] Hutchinson-Gilford progeria syndrome (HGPS), atypical Werner’s syndrome (WS), restricted dermopathy (RD) and mandibuloacral dysplasia (MAD) [7,8,9]collectively termed as laminopathies. DCM is characterised as a severe cardiac muscle disorder, resulting in ventricular dilation and impaired systolic functions. Till date, 200 *LMNA* mutations have been linked to DCM (http://www.umd.be/LMNA/) which accounts for 5%-10% of DCM cases [10,11,12]. Various studies attributed the involvement of epigenetic modifications as one of the major underlying causes of cardiovascular diseases [13,14,15]added more references.

Lamins have been shown to interact with chromatin at Lamina Associated Domains (LADs) [16] polynucleosomes, histones, and DNA [17,18,19,20]. Association of the lamins with the peripheral heterochromatin justifies its role as an epigenetic regulator [2], [21,22]. Although lamin A/C associated LADs are rich in euchromatic regions, B-type lamin associated LADs are heterochromatic in nature [23]. The LAD vary from 0.1-10Mb in length and the genes within the LADs usually remain repressed with epigenetic modifications like H3K9me2 and H3K9me3 [24,25]. In HGPS histone modification pattern is drastically modified [26,27]. On the other hand, lamin A/C deficiency has been shown to cause loss of heterochromatin and is associated with rearrangement of Heterochromatin Protein 1β (HP 1β) and H3K9 methylation [28,29]. Changes in lamin A/C chromatin interaction imply rearrangement of chromatin organization which is controlled by epigenetic processes [30]. This brings us to a juncture where the association of epigenetic changes in laminopathies becomes an important avenue to explore. Borrowing cues from our previous studies, we investigated the effect of *three* such DCM causing lamin A mutants on the alteration of the epigenetic landscape in myogenic fibroblast cell lines C2C12. We have focused on K97E, R377L, and S573L, as these mutations are reported to cause DCM in various populations with severe phenotypes [31,32,33,34]. In retrospect, the severity of phenotype caused by K97E which lies in the rod domain had been already established from detailed *ex vivo* studies [35]. R377L resides at the extreme end of the rod domain, whereas S573L lies near the Ig fold domain of lamin A (Figure 1). We showed through immunofluorescence analyses that expression of mutant lamins can alter overall localization pattern of histone modification marks within the nucleus. Expressions of the mutant lamin A proteins are also responsible for anomalous aggregation of RNA Pol II. Taken together we have established for the first time, how these mutants could potentially alter the gene expression profile which in turn might play deterministic roles in the pathogenesis of DCM.

**Figure 1:**
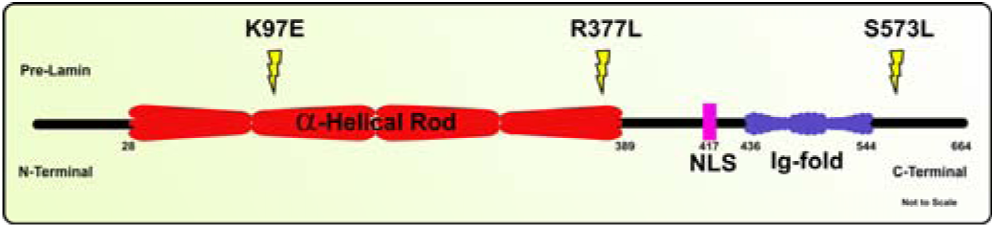
Schematic representation of position of mutants in Human full length lamin A.

## Results

### Mutant Lamin A Alters the Nuclear Distribution of Histone Modification Marks

Alteration in global histone modification marks were reported to be associated with various lamin A mutations [18,26,27,28]. To study the effects of DCM associated lamin A mutants K97E, R377L and S573L on histone modification marks, we focused on few such marks which are involved in both transcription activation, repression or silencing such as H3K4me3, H3K36me3, H3K9me3 and H3K27me3 respectively. We have ectopically expressed these mutations in C2C12 lines which is a murine myoblast cell line. As the above mentioned mutations are associated with cardiac muscle disorder, it justifies the choice of this particular cell type. Condensed heterochromatin are enriched in H3K9me3 and H3K27me3[36] whereas the marks H3K4me3 and H3K36me3 are associated with open chromatin configuration and is, therefore, characteristic of euchromatin [1,37]. As previously reported, overexpression of K97E causes abnormal nuclear lamina formation along with differently sized aggregates of lamin proteins inside the nucleus [35]. We observed aggregates of lamin in K97E expressing C2C12 cells; however the aggregates were milder in R377L expressing cells. Interestingly enough, S573L mutation showed no abnormality either in terms of lamina organization or any lamin aggregation foci (Figure 2). Our immunofluorescence analysis for the H3K9me3 marks showed colocalization of lamin aggregates in K97E and R377L mutants (Figure 2). Whereas H3K4me3 staining pattern revealed cells expressing R377L mutation, despite having milder phenotype compared to K97E, exhibited sequestration within the lamin aggregates (Figure 2). However, chromatin containing H3K27me3 marks for K97E, R377L and S573L revealed a distribution pattern similar to that of the wild type (WT) transfected cells (Figure 2). As shown in figure 2 the staining pattern of H3K4me3 also showed no abnormality for K97E and S573L mutation. H3K36me3 marks on the chromatin for all the mutants showed no anomalies (Figure 2). Thus, the distribution patter of chromatin marks containing H3K9me3 and H3K4me3 seemed alter in K97E and R377L mutants. Aggregates formed by mutant lamin A expression co-localized with the DAPI marks which stains the pericentric heterochromatin (Figure 2). H3K9me3 mark helps to recruit heterochromatin protein 1 (Hp1) to various regions of chromatin. Hp1 is known to play structural role in regulating the heterochromatin organization and gene expression [38,39,40]. Based on the changes in H3K9me3, we examined the distribution pattern of HP1γ [26,41,42]. We observed that, HP1γ was sequestered at places where lamin aggregates were present for K97E along with the accumulation of H3K9me3 marks (Figure 3). Thus, above observations indicate a global reorganization and redistribution of specific histone modification marks due to expression of mutant lamin A particularly K97E.

**Figure 2:**
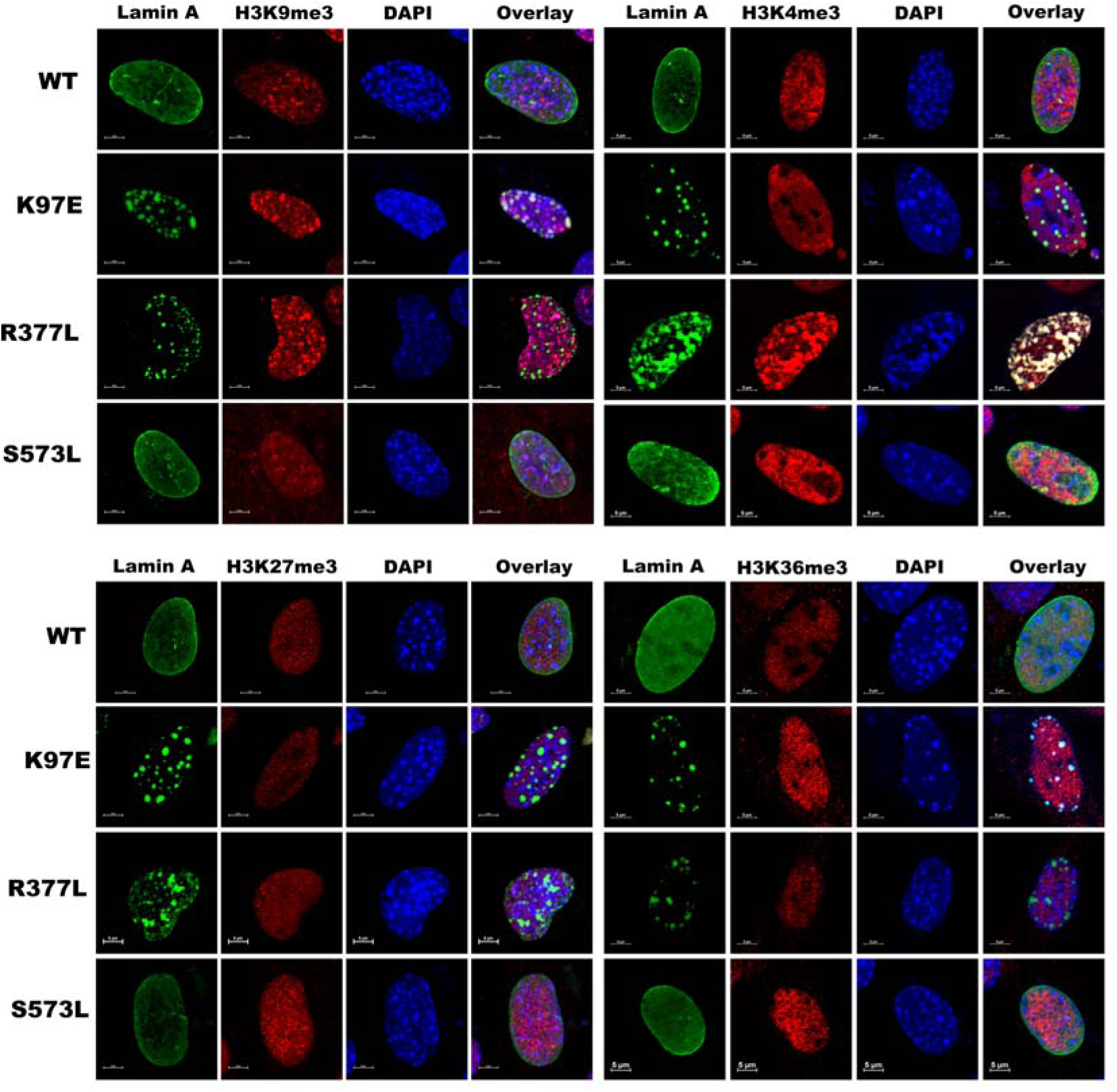
Alteration of global histone modification marks. C2C12 cells were transfected with pEGFP-human-WT, K97E, R377L and S573L – LA for 24hours and stained with anti H3K9me3, H3K27me3, H3K4me3, H3K36me3 antibodies for histone modification marks which are depicted in red and lamin A in green. Scale bar is 5μm.

**Figure 3:**
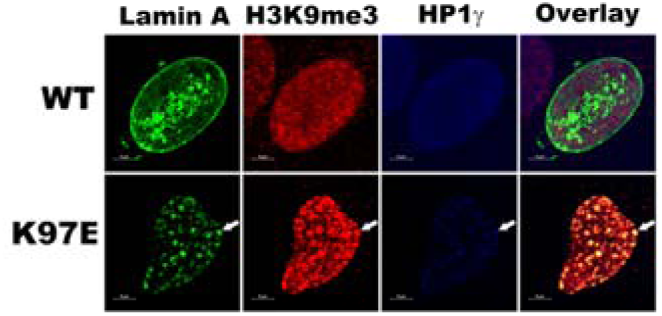
Alteration of Hp1γ in cells expressing lamin mutation. C2C12 cells were transfected with pEGFP-human-WT, K97E- LA for 24hours and stained with anti HP1γ and H3K9me3 antibodies which are shown in blue and red respectively. Scale bar is 5µm.

### Mutant Lamin A Alters RNA Pol II Distribution

Numerous experimental evidences support the concept that lamins play both direct and indirect roles in RNA Pol II mediated transcription [43]. Based on these above results we further checked the levels of activated hyperphosphorylated forms of RNA Pol II (Pol IIos2 and s5) labelling in the nuclei of cells expressing wild type and mutant lamin A as shown in Figure 4. Phosphorylation at Serine 2 of RNA Pol II (RNA Pol IIs2p) indicates transcription elongation, whereas phosphorylation at Serine 5, of RNA Pol II (RNA Pol IIs5p) indicates transcription initiation [44]. Our data showed that aggregates formed by K97E and R377L colocalized with the RNA polymerase containing the serine 2 phosphorylation. However, RNA polymerase containing the serine 5 phosphorylation was only sequestered by K97E lamin aggregates. Thus, the K97E aggregates severely affected the localization of RNA polymerase. However, R377L aggregates only seemed to alter the distribution pattern of RNA Pol IIs2p which regulates the transcription elongation.

**Figure 4:**
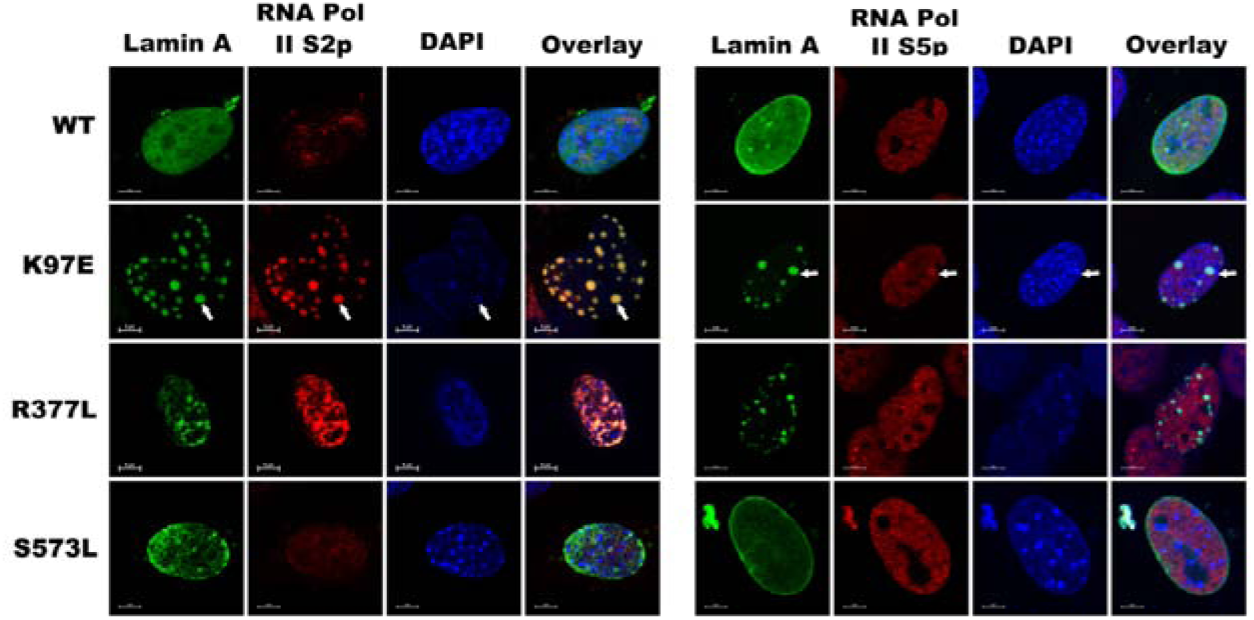
Global RNA Pol II s2p and s5p distribution. C2C12 cells were transfected with pEGFP-human-WT, K97E, R377L and S573L – LA for 24hours and stained with anti RNA Pol II s5p, RNA Pol II s2p antibodies separately and shown in red. Scale bar is 5µm.

## Discussion

An emerging hypothesis in the field regarding molecular mechanism of laminopathies states that mutations in *LMNA* gene leads to aberrant gene expression profile during differentiation in a tissue specific manner [45,46,47]. These changes in gene expression are largely controlled by epigenetic modulations in the chromosomes [48,49]. In few laminopathies, mutation or loss of lamin A/C has already been linked with alteration in chromatin organization at epigenetic levels [26,27,28,29]. These studies mostly dealt with specific laminopathies which include HGPS and muscular dystrophies. However, till date there are few studies reporting alteration in epigenetic landscape and its association with the lamin A/C mutation in DCM [50,51]. Mutations in the rod domain of lamin A account for almost 60% of the reported mutations for the laminopathies. However, the tail region of lamin accounts for 35% of the mutations associated with laminopathies (http://www.umd.be/LMNA/) [52]. Hence we decided to study the mutations K97E in rod 1B, R377L in rod 2B and S573L in the tail region. In this report the immunofluorescence data revealed that lamin A K97E led to the sequestration H3K9me3 in the lamin foci. Interestingly, R377L showed sever alteration in H3K4me3 marks distribution. This prompted us to conjecture that the function of transcription machinery might be perturbed in such a scenario. Methylation of histones is associated with either activation or repression of genes. For instance, tri-methylation at H3K9 position is associated with gene silencing whereas tri-methylation at H3K4 position is associated with gene activation [53]. Active genes possess a high enrichment of H3K4me3, which marks the transcriptional start site (TSS) [54,55]. Whereas H3K9me3 marks are reported to be associated with transcription silencing and heterochromatin formation[56]. Interestingly, these associations of mutant lamin A aggregates with epigenetic markers are not generic and different types of mutations showed specific association with specific epigenetic marks. However, lamin A S573L showed no significant alteration in its interaction with chromatin. The fact that nuclear periphery is enriched in condensed heterochromatin, justifies the association of H3K9me3 marks in K97E aggregates. Though lamins at the periphery are known to recruit condensed heterochromatin [57] however, reports have mentioned the existence histone marks for active chromatin near the nuclear pore complexes (NPC) [58]. Heterochromatin Protein 1 (HP1) recognises H3K9me3 – mark associated with repressive heterochromatin. HP1binds to H3K9me3 via its N-terminal chromo domain and this interaction is important for the overall structure of heterochromatin [41,42]. Constitutive heterochromatin is characterised by relatively high levels of H3K9me3 and HP1α/β[59] but not HP1γ which reside within euchromatin [60,61] and are known to colocalize with the coding regions of actively transcribed genes in mammalian cells, suggesting a role in active transcription [62]. Moreover a report by Scaffidi and Misteli in 2006 [26] has shown that heterochromatin markers, such as H3K9 trimethylation and heterochromatin-associated protein HP1γ, are reduced in patient cells from HGPS. Lamin A/C deficiency causes loss of heterochromatin which is also further associated with rearrangement and dispersion of HP1β and H3K9 methylation within the nucleoplasm [28,29]. This was further strengthened when we looked into the status of HP1γ in K97E expressing cells where changes in H3K9me3 marks had been observed. We noticed co-localization of HP1γ with the foci of laminA aggregates. Our results are consistent with previous reports where a variety of epigenetic histone marks are altered in HGPS cells expressing mutant lamin A, which includes reduction in the levels ofH3K9me3and increase in H4K20me3[26,27]. Therefore, we can hypothesize that lamin A does indeed modulate epigenetic landscape via interactions with different chromatin remodelling proteins.

RNA polymerase activity partially depends on its interaction with the structural components of the nucleus which happens to be lamin proteins [65]. Immunostaining of both for RNA Pol II S2p and S5p marks in K97E transfected cells showed the abnormal distribution of RNA polymerase, whereas in R377L transfected cells, we noticed accumulations of RNA Pol II S2p marks which indicate a probable elongation defect in the transcription process. Interestingly, the transcription initiation process which is marked by RNA Pol II S5p showed no anomaly in R377L mutants. This active chromatin signature in the aggregates might promote the transcription machinery binding and affect the proper functioning of the transcription machinery complex. Nonetheless any such sequestration perturbs the equilibrium distribution of the interacting proteins needed for RNA Pol II mediated transcription. Therefore it can be postulated that K97E alters the epigenetic landscape of the C2C12 cells, which in turn modifies the transcription circuitry hitherto unknown. This is the first report encompassing 3 different mutations of lamin A implicated in the same disease, where we arrived at a consensus model of differential epigenetic regulation. It is beyond the scope of the work to establish a correlation between differential gene expressions as a sequel to epigenetic modulation, which nonetheless keeps the avenue of further research open.

## Materials and Methods

### Site Directed Mutagenesis

Site directed mutagenesis was performed on pEGFP-human lamin A (gift from Dr. Robert D. Goldman, Northwestern University, Chicago)following the protocol as described in [35] to obtain the mutations K97E, R377L and S573L. Primer sets each containing the mutations as listed in table S 1.1 were used to develop the mutants. Mutants were confirmed by Sanger sequencing using the sequencing primers for SDM as listed in table S 1.2

### Cell Culture

C2C12 cells were grown and cultured in High Glucose Dulbecco’s modified Eagle medium (DMEM) (Gibco), complemented with 10% fetalbovine serum (Gibco) and 1% penicillin-streptomycin (Gibco), in a humidified incubator at 37°C in 5% CO2. Cells grown on glass coverslips (Himedia)were synchronized at G2/M phase with nocodazole (Sigma). C2C12 cells were treated with 200 ng/ml of nocodazole for 12 h [66]. Cells were washed three times with PBS and replenished with pre-warmed fresh complete media after release form the block. Cells were transfected with pEGFP-human LA (EGFP-wt LA) and mutants after 3h from release of the nocodazole block using Lipofectamine 2000 (Invitrogen) in accordance with the manufacturer’s protocol at early passages having 80-85% confluency and were kept in culture for 24 hrs [67]; the DNA/lipofectamine ratio was kept at 1:1.5. Enrichment of the EGFP positive cells ectopically expressing pEGFP-human LA (EGFP-wt LA& EGFP-mut LA), was obtained by sorting using FACS Calibur platform from BD Biosciences (California, USA) equipped with 488 nm laser system for further studies.

### Indirect Immunofluorescences

Post transfections cells were fixed with 4% paraformaldehyde and permeabilized with 0.5% Triton X-100 at RT. Thereafter, cells were incubated with primary and secondary antibody. Coverslips were mounted with Vectashield containing DAPI for staining DNA. Primary and Secondary antibodies used are described in table S1.3.

### Image Acquisition

The slides were visualized in NIKON Inverted Research Microscope ECLIPSE TiE with Plan Apo VC 100X oil DIC N2 objective/1.40 NA/1.515 RI with a digital 4X zoom. The images were captured with A1R+ confocal scanner head (Galvano mode) mounted on Ti-E inverted microscope. The excitation filters used were 1) First filter cube: 450/50, 2) Second filter cube: 525/50, 3) Third filter cube: 595/50, the first dichroic mirror used was 405/488/561. While capturing the images, pinhole was maintained at 57.5 μm, one way scan direction and channel series mode line 1->4 was used. While capturing the Z stacks a step size of 0.25 μm was maintained. The wavelengths and specifications of different lasers used were 1) CH1: Multi line Argon-Krypton mixed gas laser of 40 milli watt, λ_ex_ -457/488/514 nm. 2) Solid state laser 100 milli watt, λ_ex_ -405 nm, 3) Solid state laser 20 milli watt, λ_ex_ -561 nm. Images were processed using Ni Elements Analysis AR Ver 4.13.

## Acknowledgement

The vector pEGFP-LA was generous gift from Dr. Robert D Goldman, Feinberg School of Medicine, Northwestern University, Chicago. The authors sincerely thank Dr.Chandrima Das, Biophysics Division, SINP, for sharing her valuable antibodies during initial experimental setup. AB thanks UGC for the fellowship. KSG thanks MMDDA and BARD projects of DAE, Govt. of India.

## Author contributions

AB performed major experiments. AB and KSG wrote the manuscript. KSG conceived the entire project.

