## Supplementary Information for "Alteration of epigenetic landscape by lamin A mutations: Hallmark of Dilated Cardiomyopathy"

**Supporting Information file for “Alteration of epigenetic landscape by lamin A mutations: Hallmark of Dilated Cardiomyopathy”**

**Avinanda Banerjee1, and Kaushik Sengupta1,2***

**1)	Biophysics and Structural genomics Division, Saha Institute of Nuclear Physics, 1/AF, Bidhannagar, Kolkata-700064, India.**

**2)	Homi Bhabha National Institute, Mumbai**

***To whom correspondence should be addressed**

Key words- Dilated Cardiomyopathy, *LMNA,* epigenetics, Histone modifications, RNA Pol II, RNA sequence analysis,

*Running Title: DCM afflicted LMNA mutants alter hetero and euchromatin organization leading to differential gene expression*

***SI Methods***

***Site Directed Mutagenesis***

Table S 1.1 Primers for SDM

| **Point Mutation** | **Sequence 5’→3’** |
| --- | --- |
| **K97E** | **gactcagtagccgaggagcgcgccc** |
| **K97E_antisense** | **gggcgcgctcctcggctactgagtc** |
| **R377L** | **ctggcggaaaaggagcgggagatggccgag** |
| **R377L_antisense** | **cctccaagagcttgaggtaggcgtggatc** |
| **S573L** | **cccactgcagcagcttgggggacc** |
| **S573L_antisense** | **ggtcccccaagctgctgcagtggg** |

**Table S 1.2: Primers for Sequencing SDM**

| **Sequencing primer** | **Sequence 5’→3’** |
| --- | --- |
| **K97E Forward** | **gggctgcgcc ttcgcatcac cgagtctgaa** |
| **K97E Reverse** | **caggtcaccctccttcttggtattgcgcgc** |
| **R377L Forward** | **ctggcggaaa aggagcggga gatggccgag** |
| **R377L Reverse** | **tttggtgacgctgcccccaccctgtgtctg** |
| **S573L Forward** | **gtggccatgc gcaagctggt gcgctcagtg** |
| **S573L Reverse** | **ggcagaagagccagaggagatgggtccgcc** |

***Indirect immunofluorescence***

**Table S 1.3: List of Antibodies Used in Indirect immunofluorescence**

| **Primary Antibodies** | **Dilution** | **Secondary Antibodies** | **Dilution** |
| --- | --- | --- | --- |
| **Anti-Histone H3 (tri methyl K27) Active Motif (39155)** | 1:1500 | **Goat anti-Rabbit IgG (H+L) Secondary Antibody, Alexa Fluor® 594 conjugate Invitrogen (A11012)** | 1:1000 |
| **Anti-Histone H3 (tri methyl K36) Abcam (ab9050)** | 1:5000 | **Goat anti-Rabbit IgG (H+L) Secondary Antibody, Alexa Fluor® 594 conjugate Invitrogen (A11012)** | 1:1500 |
| **Anti-Histone H3 (tri methyl K4) Active Motif (39159)** | 1:1000 | **Goat anti-Rabbit IgG (H+L) Secondary Antibody, Alexa Fluor® 594 conjugate Invitrogen (A11012)** | 1:800 |
| **Anti-Histone H3 (tri methyl K9) Active Motif (39161)** | 1:1000 | **Goat anti-Rabbit IgG (H+L) Secondary Antibody, Alexa Fluor® 594 conjugate Invitrogen (A11012)** | 1:800 |
| **Anti-HP1 γ Millipore (05-690)** | 1:500 | **Goat anti-Mouse IgG (H+L) Secondary Antibody, Alexa Fluor® 405 conjugate Invitrogen (A31553)** | 1:600 |
| **Anti-RNA polymerase II CTD (4H8) Abcam (ab5408)** | 1:1000 | **Goat anti-Mouse IgG (H+L) Secondary Antibody, Alexa Fluor® 594 conjugate Invitrogen (A11005)** | 1:800 |
| **Anti-RNA polymerase II CTD (phospho S2) [H5] Abcam (ab24758)** | 1:800 | **Goat anti-Mouse IgG (H+L) Secondary Antibody, Alexa Fluor® 594 conjugate Invitrogen (A11005)** | 1:800 |
